# Sense of Agency for Mental Actions: Insights from a Belief-Based Action-Effect Paradigm

**DOI:** 10.1101/2021.01.30.428924

**Authors:** Edmundo Lopez-Sola, Rubén Moreno-Bote, Xerxes D. Arsiwalla

## Abstract

A substantial body of research in the past few decades has converged on the idea that the so-called “sense of agency”, the feeling of being in control of our own actions, arises from the integration of multiple sources of information at different levels. In this study, we investigated whether a measurable sense of agency can be detected for mental actions, without the contribution of motor components. We used a fake action-effect paradigm, where participants were led to think that a motor action or a particular thought could trigger a sound. Results showed that the high-level sense of agency, measured through explicit reports, was of comparable strength for motor and mental actions. The ‘intentional binding’ effect, a phenomenon typically associated with the experience of agency, was also observed for both motor and mental actions, with the only exception of short action-effect delays. Furthermore, a consistent relationship between explicit reports of agency and intentional binding was found. Taken together, our results provide novel insights into the specific role of intentional cues in instantiating a sense of agency, even in the absence of motor signals. These results may have important implications for future brain-computer interfaces as well as for the study of pathological disruptions of agency.

## 1. Introduction

“Sense of agency” (SoA) refers to the subjective experience or feeling of being in control of our actions and their consequences. When we walk, type, or perform voluntary actions, we hold a subjective feeling of being the agents initiating those actions. How the sense of control over our actions is constructed has been the subject of a large body of research in the last twenty years, and is still a highly debated problem in cognitive science [1].

Different accounts have been put forward to explain how the sense of agency is constructed. The early years saw a dichotomy between models based on postdictive inferential processes (D. Wegner’s theory of apparent mental causation [2]) and models relying on predictive mechanisms (comparator model theories [3]). This has progressively been replaced by unifying accounts that consider both types of signals as crucial for the realization of a sense of agency [4, 5, 6]. According to these views, the SoA arises from the integration of multiple cues of distinct origins, such as motor signals, sensory feedback, background beliefs and affective cues. In fact, it has been argued that different cues could be integrated at different levels. Synofzik and colleagues [6] have proposed distinguishing between a low-level, pre-reflective feeling of agency (FoA), and a high-order, explicit judgment of agency (JoA). The FoA would arise at the sensorimotor level from the interplay of predictive signals (motor predictions, efference copy) and postdictive inferences, and corresponds to the “raw” feeling of being in control of an action. The JoA refers to explicit attributions of authorship (“I did this”) and emerges at the conceptual level, predominantly based on the low-level FoA but also influenced by contextual cues and high-order beliefs.

Despite the general consensus that sense of agency depends on the integration of multiple cues at different levels, the specific contribution of each of these cues to each level is still a matter of debate [4, 6]. In particular, the relevance of motor signals has been repeatedly emphasized, and some authors have suggested that without motor predictions the SoA would be largely reduced [7, 8, 9]. However, to date most experimental paradigms do not conclusively address the issue of whether or not the SoA could emerge in the absence of motor cues. One specific limitation of most paradigms comparing the SoA for motor and non-motor actions is that the participants’ intentionality to produce an action is usually absent in the non-motor condition [8, 9]. This is a problem, since intentionality is a crucial element in the construction of the SoA. In order to dissociate the concrete role of motor predictions from other mechanisms contributing to the SoA, one would need to design a paradigm where an *intentional* overt motor action is contrasted with an *intentional* action that does not involve motor signals.

It is intuitive to think that in a situation where an intentional mental event (e.g., thinking about moving an object) would be followed by the desired outcome (the object moved), the match between intention and sensory feedback would lead to the emergence of a sense of agency. Interestingly, the use of Brain-Computer Interfaces (BCI) allows subjects to produce controlled outcomes in the outside world, such as moving a prosthetic arm or controlling a cursor on a computer screen, through neuro-psychological processes that do not necessarily involve bodily movements [10]. In the following, we will refer to such neuro-psychological processes as *mental actions*. The term *mental action* has been intensively discussed in the philosophy of mind and action, and it is unclear which specific mental events should be regarded as mental actions [11, 12]. Here, we adopt the term *mental action* for that in which the cognitive capacities of an agent are recruited to produce an effect of some kind, for example, “solving a chess problem in one’s head, or deliberating about whether to accept a job offer” [13]. The mental action thus defined can eventually trigger outcomes in the external world through the interaction with BCIs (such actions are also referred to as “BCI-mediated actions”).

Several studies have investigated the SoA arising from the interaction with BCIs [14, 15, 16, 17, 18, 19] most of them suggesting that a SoA may result from BCI-mediated actions under appropriate conditions. This is coherent with the aforementioned idea that intentional mental actions could lead to a subjective experience of control without the presence of motor commands. All these studies have assessed the SoA with explicit reports of agency, and therefore, they have so far only measured high-level JoA from participants.

At this stage, several questions remain unanswered: First of all, it is unclear if the SoA resulting from mental actions is comparable to the strong experience of agency resulting from motor actions. Secondly, if mental actions do lead to a SoA, as indicated by BCI studies, it is not known whether this SoA emerges only at a conceptual level (JoA) from the combination of thoughts, intentions and contextual cues, or if a low-level FoA also exists at the non-conceptual level in the absence of motor signals.

In the present study, we have investigated these issues by contrasting the SoA arising from motor actions with the one resulting from mental actions. To that purpose, we have used an action-effect paradigm where a given motor or mental action was followed by a sound. In the motor action condition, participants had to press a key. In the mental action condition, a “fake” BCI was used: participants were led to believe that one specific thought could be detected by a computer software (see “Methods”), which would subsequently produce a sound. Other studies have succeeded at creating an illusion of control using fake BCIs [17, 16]. The paradigm is based on the idea that a causal action-effect link is not necessary for the emergence of SoA. It is sufficient that the action and its alleged effects are close enough in time and that the subject *believes* there exists a causal relationship between both events. Importantly, in order to ensure that both experimental conditions were adequately similar in this study, no causal relationship existed in the motor action condition: participants were led to think that their key press would eventually produce a sound, but the sound always appeared independently of the participants’ action.

Based on previous studies [20, 9, 21], we manipulated the time interval between the action and the effect. Temporal contiguity between an action and its outcome is a strong indicator of authorship, and shorter intervals typically lead to a stronger SoA [22, 21]. Additionally, in order to introduce a certain degree of ambiguity in the attribution of authorship, participants were told that the sound could be produced either by their action or by the computer.

We used explicit and implicit measures of the sense of agency [1]. As an explicit measure, we asked participants to report whether they had produced the outcome or not (authorship report). The implicit measure used was the intentional binding (IB) paradigm [23]. In their seminal study, Haggard and colleagues found that time perception was altered upon execution of voluntary actions: the action, e.g. a key press, was perceived later in time, and its effect, e.g. a sound, was perceived as occurring earlier; so that both events were subjectively perceived to be closer in time. Crucially, this effect was not found when the action was involuntary (a TMS pulse over the brain motor area was applied to contract the muscle that pressed the key). We used both types of measures to clarify at which level the SoA for mental actions is constructed, explicit reports being a marker of high-level agency, and the intentional binding effect being typically associated with a low-level perceptual feeling of agency [24].

The main hypothesis of this study is that mental actions can produce a sense of agency, and that this sense of agency will be comparable to that of motor actions. In particular, we hypothesize that:

- The judgement of agency (JoA), measured at the conceptual level by explicit reports, will be similar for mental actions to the one for motor actions. We predict that for shorter action-effect intervals participants will attribute the sensory effect more frequently to themselves than to the computer; and that this frequency will be comparable for motor and for mental actions.
- Regarding the low-level (perceptual) feeling of agency (FoA), some authors have recently argued that predictive mechanisms based on counterfactuals and high-order cognitive sources (intentions, beliefs, etc.), different from predictive motor models, could play a key role in several body-related action-outcomes, including agency attribution [25]. Based on this idea, and on the fact that other mechanisms such as inferential postdictive processes may also contribute to the feeling of agency [6], we hypothesize that a low-level FoA will arise from mental actions. This is of special interest for the current research on agency, since, to our knowledge, this is the first study investigating low-level agency in the absence of motor cues. Our prediction is that mental actions will produce intentional binding to an extent similar to that of motor actions. It is worth noting that some studies have suggested that the IB effect only arises when motor cues are present [9], while others have argued that it is a general signature of intentionality, independent of motor commands [26, 27]. Others have even claimed that it reflects the mere experience of causality [28]. The present study seeks to clarify whether the IB effect is specifically linked to motor actions or not.
- Finally, given that a higher sense of agency is expected for short action-effect delays, our prediction is that the IB effect will be found for short, but not for long intervals, and that this will be the case both for motor and mental actions. However, some studies using the IB paradigm have not found different binding for different time intervals [24], and the relationship between IB and explicit measures of agency is currently unclear [29, 30, 31]. This study will offer a novel insight into the relationship between implicit and explicit measures of agency, both with and without motor cues.

Critically, these predictions are valid only for participants that actually believe that their motor or mental actions are causally linked to the sound (in reality, this causal link does not exist). Therefore, it is crucial for the experiment that most participants feel that their actions can produce the alleged effect. In this sense, we expect more participants to have causal beliefs for the motor condition than for the mental condition, given the increased difficulty of creating causal beliefs for mental actions (see “Methods”).

## 2. Methods

It is worth mentioning that the experiment was initially thought to be run in a laboratory environment. However, due to the SARS-CoV-2 pandemic, the experiment was adapted to be performed online. More details will be given in the following paragraphs.

### 2.1. Participant selection

50 participants (23 male, 22 female, 5 unknown) were recruited through the online platform Prolific (https://www.prolific.co/). To ensure that the instructions of the experiment were correctly understood by the participants, only English native speakers were selected.

Seven participants were excluded from the study due to system incompatibilities and one participant was excluded due to poor performance in the tasks.

### 2.2. Experimental design

#### 2.2.1. Software tools

The study was designed with the open-source python-based software Psychopy3 [32]. The experiment was translated into JavaScript and uploaded to the online platform Pavlovia (https://pavlovia.org/). The study URL from Pavlovia was provided to the participants for the execution of the experiment in their own devices. The full python and Javascript codes for the study are available at https://gitlab.pavlovia.org/edmundolopez/agency_exp.

#### 2.2.2. “Fake” BCI: a facial motion detection software

For the experiment to be successful, it was crucial to ensure that participants believed that their mental actions could trigger an external event through the apparent BCI. Participants were told that they would participate in an experiment to evaluate the users’ interaction with a facial motion detection software. They were told that this software could detect micro-gestures in the faces of participants, which were recorded with their device webcam. They were informed that the software could recognize involuntary micro-expressions in their faces when thinking about a word related to a face gesture. In this case the word was ‘Smile’. During the trials, participants had to think about the word ‘Smile’ at a particular moment and they were told that this thought would produce a micro-gesture in their face that would be detected by the software which would eventually produce a sound. Thus, they would produce a sensory event with a mental action. Critically, participants were instructed to avoid producing any voluntary face movement during the trial.

### 2.3. Experimental procedure

Participants were first given a brief introduction to the alleged purpose of the experiment: assessing user interactions with a facial motion detection software. They were told that the main objective of the study was “the sense of control and the sense of time while interacting with the software”.

The experiment consisted of three distinct blocks, one per experimental condition: first, the SoA of the subjects was assessed through explicit and implicit measures in a motor action task; then, the same measures were used to evaluate the SoA in a mental action task; finally, in the last block we measured the time estimation error made by participants when not performing any action (baseline condition).

#### 2.3.1. Motor action condition

Participants were told that in the first block of the experiment their sense of control and time over motor actions would be measured, without using the facial detection software. The motor action condition was divided in two parts: in the first part, participants performed three training trials and twelve test trials, and after each trial the sense of agency was measured via explicit report. The second part had the same number of trials, and after each trial the intentional binding was measured instead.

The design of the trials was based on the work of Haggard and colleagues on intentional binding [23]. In each trial, participants were shown a clock and a white dot rotating with a period of 2.55 seconds. A red cross appeared after 500 ms in a given position of the clock (which was in fact the position from where the white dot started rotating), and remained on screen during 500 ms, after which it disappeared. Participants were instructed to press the ‘space’ key when the white dot reached the position where the red cross had appeared. Crucially, the time when the dot reached that position was always the same (2.55 seconds after dot motion onset), so that the sound could be programmed to occur at varying intervals from the moment in which participants performed the action (if following the instructions properly). The red cross appeared at a different location each time, chosen randomly among the sixty ‘minute’ positions of the clock, so that participants could not habituate to the clock position where the key press would be made. Participants could nevertheless habituate to the 2.55 seconds period, and solve the task by learning the period and not looking at the dot. However, this possibility was deemed to be fairly remote given the difficulty of the task, and it was assumed that participants would need to track the dot rotation to perform the task. The trial structure is shown in Figure 1.

**Figure 1:**
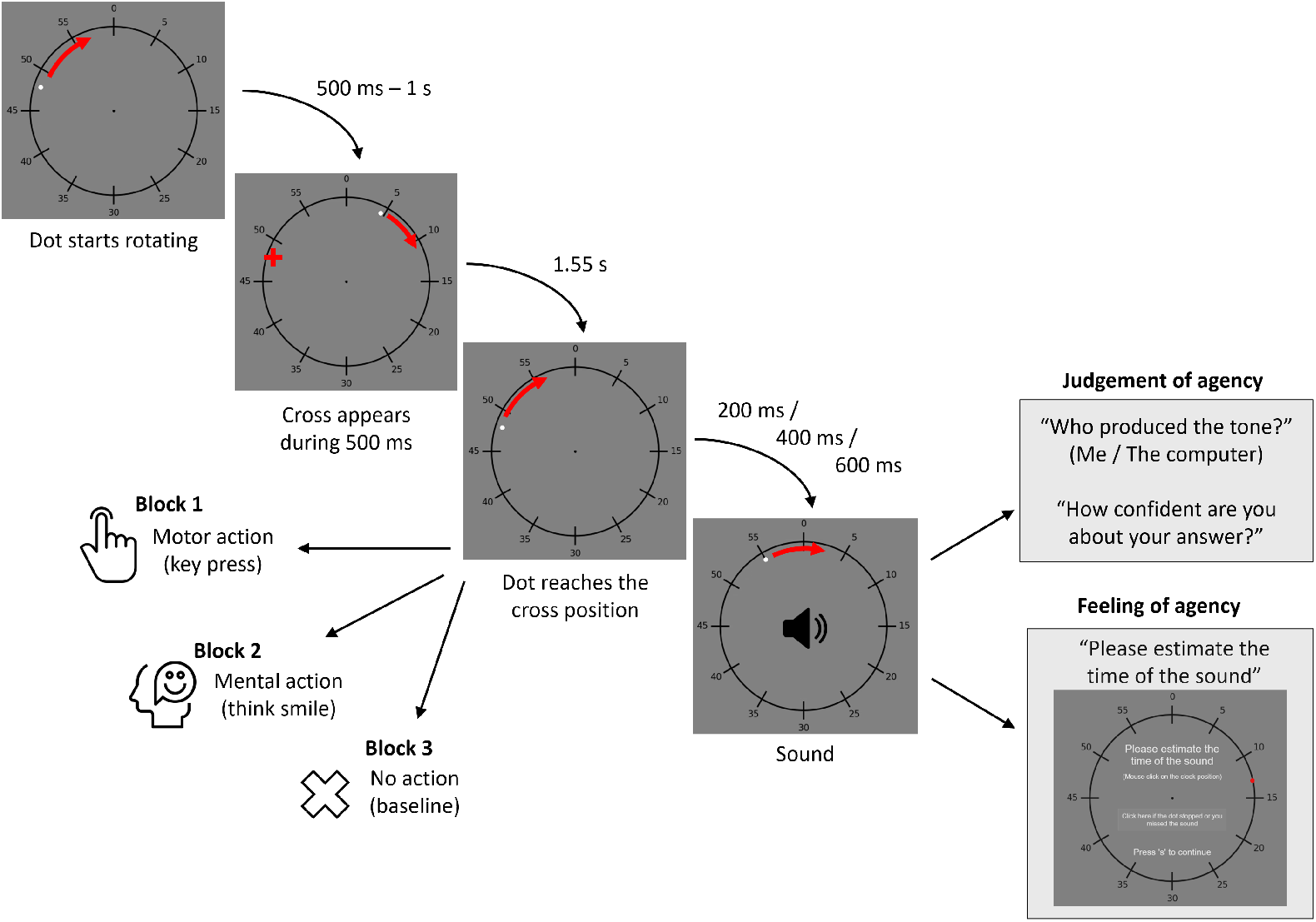
Trial structure. The dot started rotating from a random position of the clock (one out of the sixty ‘minute’ positions). After 500 ms, a red cross appeared on screen, and remained for 500 ms. The red cross appears magnified for visualization purposes. The red cross appeared at the position from where the dot had started rotating, so that the dot reached the cross position always at the same time, 2.55 ms (the dot rotation period). Participants were instructed to perform an action when the dot reached the position cued by the cross: a motor action in the first experimental block, a mental action in the second one, and no action in the third. A sound appeared after a given time interval (200, 400 or 600 ms, in randomized order) after the dot reached the cued position. After the trial, participants had to either report their judgement of agency or to judge the time when they heard the sound. It is worth noting that the stimulus presented was identical in all trials of the three blocks of the experiment, with the only difference that in each block participants were instructed to perform different actions at the cued time.

In every trial, the sound always appeared, regardless of when and whether the ‘space’ key was pressed. The sound appeared at one of three different intervals from the cued time: 200, 400 or 600 ms. Each interval appeared in total of four trials, and the order of appearance of each interval was randomized. Before the trials, participants were told that their key press would trigger a sound, but that in some of the trials the sound would be produced by the computer instead.

In the first part of this block, the JoA was assessed with explicit measures. Following each trial, participants were asked the question: “Who produced the tone?”. They could choose between “Me” and “The computer”. After this response, they had to respond using a Likert 1-7 scale to the following question: “How confident are you about your answer?”. The scale had in its extremes the labels “Not at all” and “Very much”. In the second part of the motor action condition, the FoA was evaluated. Participants had to indicate after each trial the clock position corresponding to the instant where they heard the sound, which allowed us to measure the intentional binding over the sound.

An important difference with other studies using similar paradigms to measure intentional binding is that in our study participants were instructed at which specific moment the mental or motor action had to be performed. The reason for this was that the experimental design required the subjects’ actions not to be causally linked with the sound. By instructing the participants to perform the action at a specific moment, the sound could be programmed to occur after that time (thus, after the action), thereby creating an illusion of causality. In brief, we assumed that if participants pressed the key at a given time and the sound appeared right after their action, they would feel that they produced the sound, even if no actual causal relation existed between both events and even if they were “forced” to perform the action at an indicated time.

In our experiment, only tone binding was measured, due to the fact that action binding cannot be measured in the mental action condition as there is no manner in which the actual time of the thought can be obtained. In any case, it was expected that the execution of voluntary actions (mental or motor) led to intentional binding for the sound, as has been indicated by several studies [24, 33]. In fact, it has been proposed that action binding and tone binding are driven by distinct mechanisms, and that they should be measured separately [33, 34]

#### 2.3.2. Mental action condition

Participants then completed the second block of the experiment, where their SoA for mental actions was assessed. Before the real trials, several instructions about the (fake) facial motion detection software were given to the participants. Again, the purpose of these instructions was to increase the chance that participants believed that the software could detect their thoughts through their facial micro-expressions and subsequently produce a sound, thereby creating an illusion of causality between their mental actions and the sensory outcome.

Participants completed two parts of the experiment, each consisting of one training trial and twelve test trials. As in the motor action condition, in the first part the JoA was measured by explicit report; in the second part, the FoA was measured by the intentional binding effect. Before each single trial, participants were told to position their face again for the software to operate properly. During each trial, participants viewed the clock and the rotating dot, a cross appeared in a given position after 500 ms, and they were instructed to think about the word ‘Smile’ when the dot reached the position where the cross had appeared. Again, the sound appeared at varying intervals from the cued time (200, 400, 600 ms, four times each in a randomized order). If participants performed the task correctly, they should have perceived that their thought was followed by the sound.

#### 2.3.3. Baseline condition

This last block consisted of three training trials and twelve test trials. In each trial, participants viewed the clock and the rotating dot, and a cross appearing after one second, but they were instructed not to perform any action. They were told that their webcam was not recording and that their key press would not have any effect. The sound appeared at different intervals from the moment when the dot reached the position where the cross had appeared (200, 400 and 600 ms, four times each in a randomized order).

After each trial, participants were asked to indicate in the clock the position where they heard the sound, so that the baseline error in the estimation of the sound time could be measured. This metric is required to compute the intentional binding effect: the estimated time of the sound under voluntary actions must be compared with the estimated time of the sound in a baseline non-active condition [30, 23].

#### 2.3.4. Post-experimental questions and debriefing

Once the subjects completed all three experimental conditions, they were asked two questions regarding their experience of control over the different tasks:

- “In the first block of the experiment (press key - sound), do you think that in some trials the sound was produced by your key press?”
- “In the second block of the experiment (think ‘SMILE’ - sound), do you think that in some trials the sound was produced by your thought?”

The possible answers where “Yes” and “No”. These questions are crucial for the study because they provide information about whether participants had believed or not that they could eventually produce the sound with their motor or mental actions. If a participant replied ‘Yes’ to the first or second question, this means that they thought that at least in one trial they produced the sound; therefore, the participant believed the “story” about the facial motion detection software, or about their key press producing the sound. It is critical to separate participants who had causal beliefs from those who did not because a strong sense of agency was expected only in those subjects who believed that their motor or mental actions could trigger the sound at all.

At the end of the experiment, participants were informed about the real goal of the experiment. They were told that there was no real facial motion detection software and no causal relation between their key press and the sound.

### 2.4. Data analysis

Analysis of the data was carried out in Python using the *statsmodels* Python module for statistical models [35] and the *SciPy* Python ecosystem [36].

## 3. Results

### 3.1. Post-experimental questionnaire

Out of 42 participants, 32 (76%) reported that they thought that in some trials their key press produced the sound, and 15 participants (36%) reported that they thought that in some trials their mental action (thinking ‘Smile’) produced the sound. All participants that believed in a causal relationship between their thought and the sound also believed in a causal relationship between their key press and the sound. The results shown in Sections 3.2, 3.3 and 3.4 correspond to the data of those participants who reported believing that their actions could produce the tone: 32 participants for the motor action condition, and of 15 participants for the mental action condition.

Only data of the aforementioned participants was used because it was expected that if a given subject didn’t believe that their actions could produce the tone, the sense of agency would not arise. Analyzing the data of subjects who didn’t have causal beliefs could be misleading because some subjects could have thought that there was no causal connection from the beginning, while others could have realized about that during the experiment. A large number of participants (50) was initially selected so that the number of participants for whom causal beliefs were reported was high enough.

To verify the fact that the sense of agency would not be produced (or in a very reduced manner) for participants without causal beliefs, we also analyzed the results for the ten participants who reported believing that neither their motor actions nor their mental actions could produce the tone. These results are summarized in Section 3.5.

### 3.2. Behavioral performance

Participants’ performance in the motor action condition was evaluated by measuring their key press time and comparing it with the cued time (the instant when the rotating dot reached the position where the cross had appeared). The mean error in the key press across participants was 96 ms (STD = 122 ms), which means that in general participants pressed the space bar after the cued time but not much later, and that this pattern was consistent across participants. No key press was detected in 13 trials (out of the 763 total trials in the motor action condition). Those trials were removed. Trials with outlier key press times (±3 STD of the group mean) were also removed (11 trials, less than 2% of the total).

In the mental action task, behavioral performance could not be measured, as it was not possible to know, by its own nature, the time when participants thought about the word ‘smile’. Therefore, we had to expect that participants performed the task correctly, and we could only encourage them to do so by reminding them the instructions of the task and emphasizing the role of the webcam and the facial motion detection software.

### 3.3. Explicit reports: a similar judgement of agency for motor and mental actions

In order to verify the hypothesis that the JoA would arise for both motor actions and mental actions, we analyzed if the number of self-attributions (responding ‘Me’ to the question “Who produced the tone?”) compared to the number of attributions to the computer was modulated by the condition (motor action versus mental action) and also by the action-effect delay.

We used the data of all trials where an explicit attribution of agency was reported. In 10 trials no report was given, and in two trials participants reported not having heard the sound. In total, data from 543 trials was available (approximately four trials per participant in each condition and interval).

We ran a binomial logistic regression where the dependent variable was authorship attribution (“Me”/“The computer”), and the main predictors were condition (motor/mental action), time interval (200/400/600 ms), and their two-way interaction. We found a significant main effect of time interval (*z* = −10.026, *p* < 0.0001), no effect of condition (*z* = −1.632, *p* = 0.103) and no interaction between condition and time interval (*z* = 1.573, *p* = 0.116). Post-hoc Chi-Square tests revealed that the number of agency self-attributions was significantly larger for short intervals than for medium intervals (*χ*^2^ = 83.7, dof = 1, *p* < 0.0001), and was also significantly larger for medium intervals than for long intervals (*χ*^2^ = 24.2, dof = 1, *p* < 0.0001). All pairwise comparisons were Bonferroni-corrected for multiple comparisons. To further assess whether there was a difference in the authorship attribution for the motor and mental conditions, we ran separate Chi-Square tests for each time interval. No significant difference was found for any of the time intervals (200 ms, *χ*^2^ = 2.89, dof = 1, *p* = 0.089; 400 ms, *χ*^2^ = 0.34, dof = 1, *p* = 0.558; 600 ms, *χ*^2^ = 0.1, dof = 1, *p* = 0.752). Figure 2a shows the proportion of agency self-attributions for the different time intervals and for the two experimental conditions.

**Figure 2:**
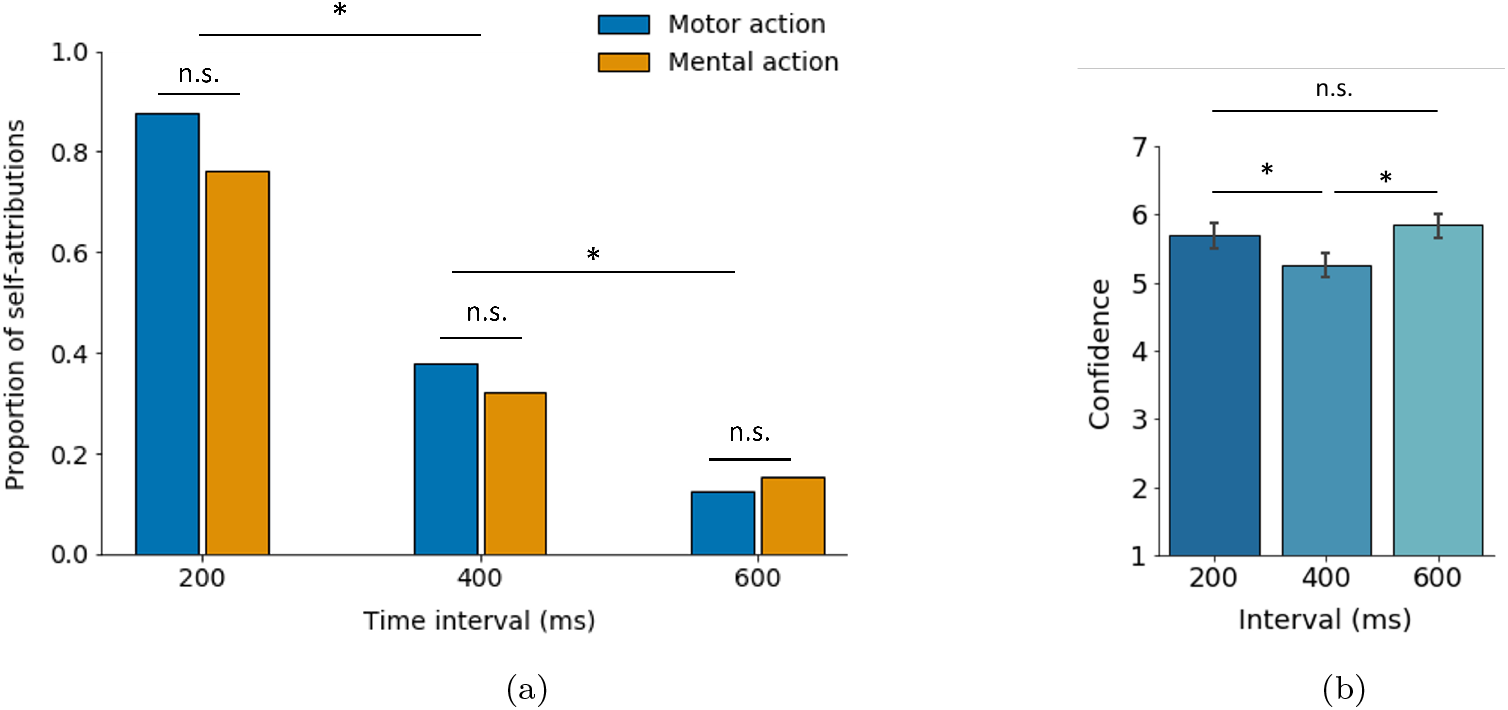
Self-attribution decreases for increased action-effect intervals but is independent of whether the action is mental or an overt motor movement. (a) Proportion of self-attributions for each condition, groups divided by interval. The statistical significance in the comparison between groups is shown for illustrative purposes (the statistical tests have not been carried out on the proportion of self-attributions but on the number of attributions). A significant effect of time interval was found, shorter intervals leading to a higher number of self-attributions. No effect of condition and no interaction were found. \**p* < 0.0001; ‘n.s.’: non-significant (*p* > 0.017); Chi-Square tests, Bonferroni corrected. (b) Mean confidence rating for the attribution of agency divided by time interval, reported in a 1–7 Likert-scale. As expected, the confidence for medium intervals is significantly lower than for short and long intervals. Error bars represent 95% confidence intervals. \**p* < 0.05; ‘n.s.’ = non-significant (*p* > 0.05); Tukey HSD test on pairwise comparisons.

Summarizing, participants thought that their action had produced the sound much more frequently when the interval between their action and the sound was short than when it was long. These results are in line with the prediction that the JoA is modulated by the time interval between the action and the sound. Crucially, there was no significant difference in the amount of self-attributions between motor and mental actions, indicating that the high-level sense of agency might have similar or homologous origins for both types of actions and is therefore observed in a similar manner in the presence or absence of motor commands.

The confidence reported by participants for their authorship attributions was also analyzed. The mean confidence rating was 5.6 (STD = 1.3) in a 1–7 Likert scale ranging from ‘Not at all’ to ‘Very much’. A between-participants two-way ANOVA on confidence ratings with time interval and action type as factors revealed a significant effect of time interval (*F*(2, 532) = 10.35, *p* < 0.0001), no significant effect of action type (*F*(1, 532) = 3.65, *p* = 0.056), and no interaction (*F*(2, 532) = 1.23, *p* = 0.292). It is a matter of discussion whether the results of Likert scale ratings can be treated as continuous variables [37]. We have done so in the present study given that this assumption is widely used, specially for 1–7 scales. A Tukey HSD test showed that the confidence rating for medium intervals (400 ms) was significantly smaller than for short intervals (200 ms, *p* < 0.05) or for long intervals (600 ms, *p* < 0.05). No significant difference was found between the confidence for short and long intervals (*p* = 0.51). This is an expected result because participants are more likely to think that they caused the sound for short intervals, and that the computer did so for long intervals, but they are less confident in attributing the authorship of the sound to themselves or to the computer when an intermediate interval is used. Figure 2b summarizes the confidence ratings for the different time intervals.

All of the analyses carried out in this section were between participants. The reason for that is that we had a different number of subjects for the action and the motor condition (see Section 3.1).

### 3.4. Implicit measurements: intentional binding appears under motor and mental actions

We analyzed the participants’ time estimations under three conditions: motor action, mental action and baseline (no action). In four trials, participants reported not to have heard the sound and in 20 trials no time estimation was given. We removed outlier time estimations, which correspond to estimations beyond ±2.5 STD of the group mean time estimation (in average less than one trial per subject). The total number of trials was 884, approximately three trials per participant per interval per condition.

To evaluate whether intentional binding was found in the motor action and mental action conditions, we compared the error in sound time estimations under each of these conditions with the error in the baseline condition. The time estimation error is calculated by subtracting the actual time of the sound from its estimated time in a given trial. For each subject, we calculated the mean baseline error and the mean error under the motor or the mental action condition. Figure 3a shows the average estimation error for all three experimental conditions. Negative errors indicate that the sound was perceived earlier than it actually occurred, and an error significantly more negative than the baseline error reflects intentional binding.

**Figure 3:**
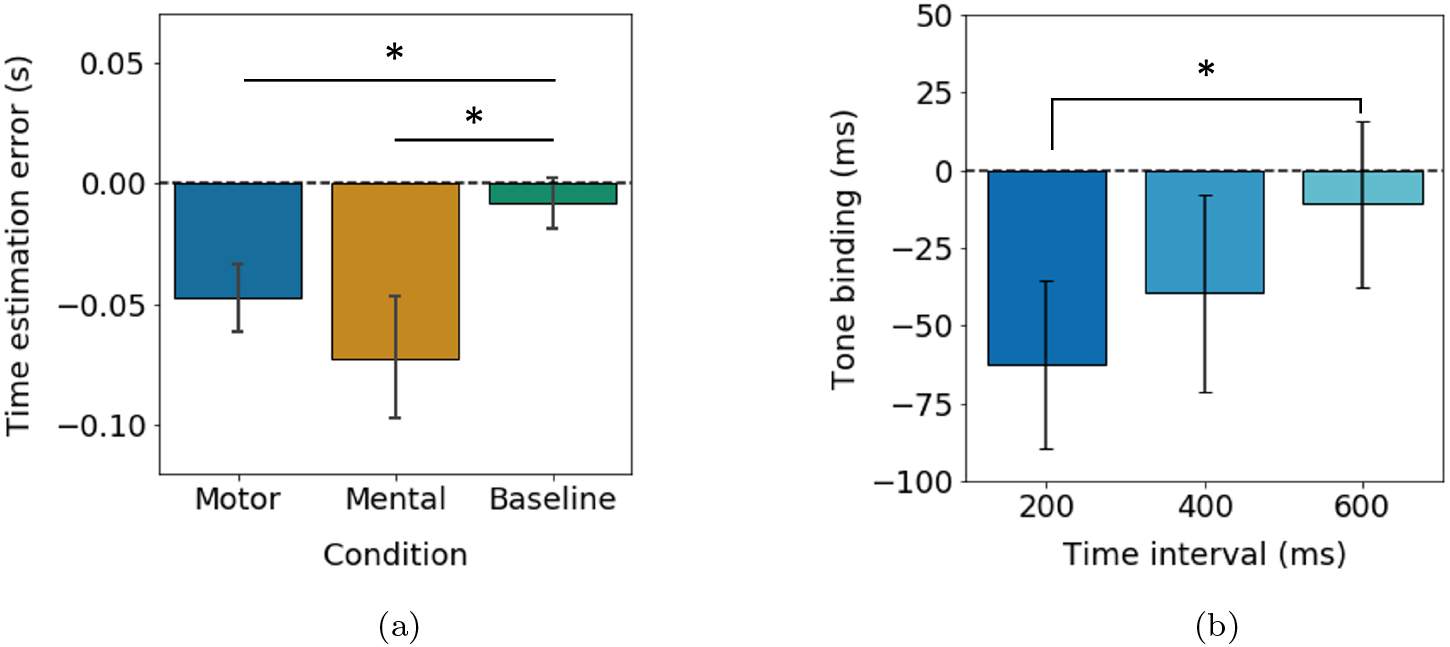
Mental and motor actions produce intentional binding over the sound, and this binding is stronger for short than for long delays. (a) Mean error in the estimation of the sound time (with respect to the actual time) for each experimental condition. Errors more negative than the baseline error indicate intentional binding. Significant binding was found for both the motor and mental action condition, indicating the presence of low-level agency. Bars represent 95% confidence intervals. \**p* < 0.05, paired sample t-tests. (b) Average binding scores for the three time delays used in the experiment, calculated as the mean estimation error of each participant in the action conditions (mental or motor) minus the mean baseline error of that participant. Shorter delays lead to higher binding, in consonance with the explicit measures of agency. Bars represent 95% confidence intervals. \**p* < 0.017, paired sample t-test, Bonferroni corrected.

Paired samples t-tests revealed that the sound was significantly anticipated in the motor action condition compared to the baseline condition (*t*(30) = −3.55, *p* < 0.05), providing evidence that intentional binding was present under motor actions. Critically, we also found that there was significant binding in the mental action condition (*t*(14) = −2.54, *p* < 0.05), which confirms the prediction that a low-level FoA may arise from non-motor actions. Independent samples t-test showed that there was no significant difference between the estimation error in the motor action and the mental action condition (*t*(44) = 0.68, *p* = 0.497), suggesting that intentional binding for the sound was produced in the same manner for both types of actions. We further confirmed that the difference in the estimation error between the two conditions was not significant with a paired sample t-test using only subjects for which data was available in both conditions (*t*(13) = 0.49, *p* = 0.636).

The previous results correspond to time estimations for collapsed intervals. Although collapsing the data for all intervals is a common way to study intentional binding [24, 9], in this study we also wanted to assess if the time interval played a role in the intentional binding effect (as it did for the higher-level SoA). First, we conducted a repeated measures one-way ANOVA on time estimation errors in the baseline condition with time interval as factor. As expected, no significant effect of time interval was found (*F*(2, 62) = 1.5, *p* = 0.231), suggesting that in the baseline condition participants did similar estimation errors for the three time intervals. Therefore, we averaged the baseline error for each participant over the three intervals and subtracted it from their estimation errors in the other two conditions, thus obtaining a direct measure of binding.

A repeated measures one-way ANOVA on sound binding scores (calculated as the mean time estimation error in the action conditions minus the mean time estimation error in the baseline condition for each participant) with time interval as factor revealed a significant main effect of time interval (*F*(2, 62) = 3.87, *p* < 0.05). Post-hoc pairwise paired t-tests showed that the binding effect was significantly larger for short intervals than for long intervals (*t*(29) = −3.31, *p* < 0.017). The significance level was Bonferroni-corrected to 0.017 for multiple comparisons. This confirms the prediction that a stronger binding would appear for short intervals, in line with the results of the explicit measures of SoA summarized in Section 3.3. The binding effect was also larger for short than for medium intervals, and for medium than for long intervals, not significantly though (*t*(29) = −1.34, *p* = 0.19; *t*(29) = −1.94, *p* = 0.06 respectively). Figure 3b displays the binding measured for the action conditions (motor and mental together) for all three time intervals.

Finally, we wanted to evaluate if, for each time interval, there was a different binding effect for the motor and for the mental actions. This allowed us to assess the relationship between implicit and explicit measures of the sense of agency: we could see if the pattern of results observed in the explicit reports (high agency for short intervals, low agency for long intervals, and no difference between motor and mental actions) is also found for intentional binding. For each time interval, we calculated every subject’s mean estimation error in each of the three experimental conditions (motor action, mental action, baseline). Figure 4 shows the average estimation error for all three experimental conditions in each time interval. For each interval, we conducted pairwise t-tests to compare the estimation errors in each action condition (motor and mental) with the baseline error.

**Figure 4:**
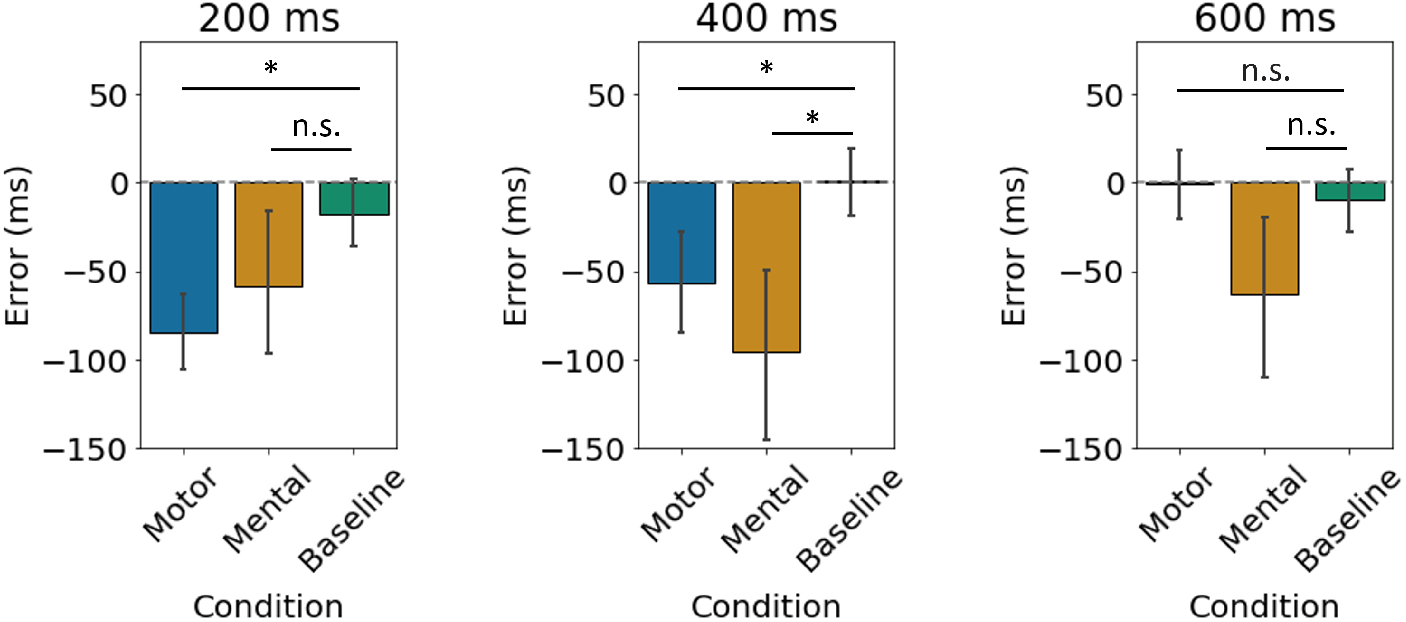
Comparison of the sound time estimation error for the three experimental conditions, data divided by time interval. For short intervals, significant binding (difference with baseline error) was found only for motor actions. For medium intervals, the binding over the sound was significant for both motor and mental actions, in line with the JoA results. For long intervals, the binding was not significant for any of these two conditions, also in consonance with the explicit reports. Bars represent 95% confidence intervals. \**p* < 0.01, ‘n.s.’: non-significant, paired samples t-test.

For short intervals (200 ms), the sound was significantly anticipated in the motor action with respect to the baseline (*t*(30) = −5.66, *p* < 0.0001), but there was no significant difference between the mental action and the baseline (*t*(14) = −0.71, *p* = 0.487). Therefore, for short action-effect delays motor actions lead to intentional binding over their effects, but mental actions do not seem to produce binding. However, an independent samples t-test suggested that the difference in time estimation errors between the motor and mental condition is not significant (*t*(45) = −1.21, *p* = 0.228), while a paired sample t-test on the subject data-set that had causal beliefs in both the motor and the action condition (14 subjects) revealed a marginally significant difference in time estimation errors in the two conditions (*t*(13) = −2.27, *p* = 0.04). Given the apparently contradictory nature of this result (marginally significant or non-significant test outcomes), it must be interpreted with caution, and it will be further elaborated in the “Discussion” section.

For medium intervals (400 ms), we found that the sound was significantly anticipated with respect to the baseline for both the motor action condition (*t*(30) = −3.32, *p* < 0.001), and the mental action condition (*t*(14) = −4.44, *p* < 0.0001). This suggests that for medium intervals binding over the tone was produced by both motor and mental actions, as predicted. Furthermore, no significant difference was found between the time estimation errors in both conditions, as shown by an independent samples t-test (*t*(45) = 1.42, *p* = 0.157) and by a paired t-test on the reduced 14 subject data-set (see above, *t*(13) = 1.03, *p* = 0.323).

For long intervals (600 ms), there was no significant difference in time estimation errors between the motor action and baseline condition (*t*(28) = 0.23, *p* = 0.820), nor between the mental action and baseline condition (*t*(13) = −1.59, *p* = 0.135). Thus, for long intervals, no binding over the tone appeared, in line with the fact that participants attributed the authorship of the sound to the computer in most of the long interval trials. A significant difference was found between the time estimation errors in the action and the motor conditions, as revealed by an independent samples t-test (*t*(45) = 2.80, *p* < 0.01) and by a paired t-test on the 14 subject data-set (*t*(12) = 2.21, *p* = 0.048). This difference suggests that participants made higher estimation errors in the mental action condition, but this is not of special relevance for the present analysis given that in none of the conditions we found significant binding.

### 3.5. “Non-believers”: participants who reported no causal beliefs displayed a reduced sense of agency

We analyzed the responses of participants who answered “No” to the two questions of the post-experimental questionnaire regarding their belief of causality in the motor and mental action conditions (10 subjects). We assume that those participants did not believe that their actions triggered the alleged effect, and that either this belief was present from the beginning of the experiment or that it appeared during the experimental tasks.

We assessed whether the explicit reports of agency in the motor and mental conditions were different for “non-believers” compared to participants who reported causal beliefs. We ran Chi-square tests to compare the number of self-attributions in each condition and each time interval for both groups. We found that, for short intervals, the number of self-attributions was significantly larger for believers than for non-believers, and that this was the case for the motor action condition (*χ*^2^ = 12.6, dof = 1, *p* < 0.001) and the mental action condition (*χ*^2^ = 9.43, dof = 1, *p* < 0.01). For medium and long intervals, no significant difference was found between believers and non-believers in none of the conditions (400 ms – motor action: *χ*^2^ = 1.76, dof = 1, *p* = 0.185, mental action: *χ*^2^ = 0.04, dof = 1, *p* = 0.841; 600 ms – motor action: *χ*^2^ = 0.13, dof = 1, *p* = 0.720, mental action: *χ*^2^ = 0.008, dof = 1, *p* = 0.927). For medium and long intervals, non-believers attributed the sound to the computer in most occasions, as was the case for participants with causal beliefs. These results, summarized in Figure 5a, confirm that the absence of causal beliefs led to a reduced (or even absent) high-level sense of agency.

**Figure 5:**
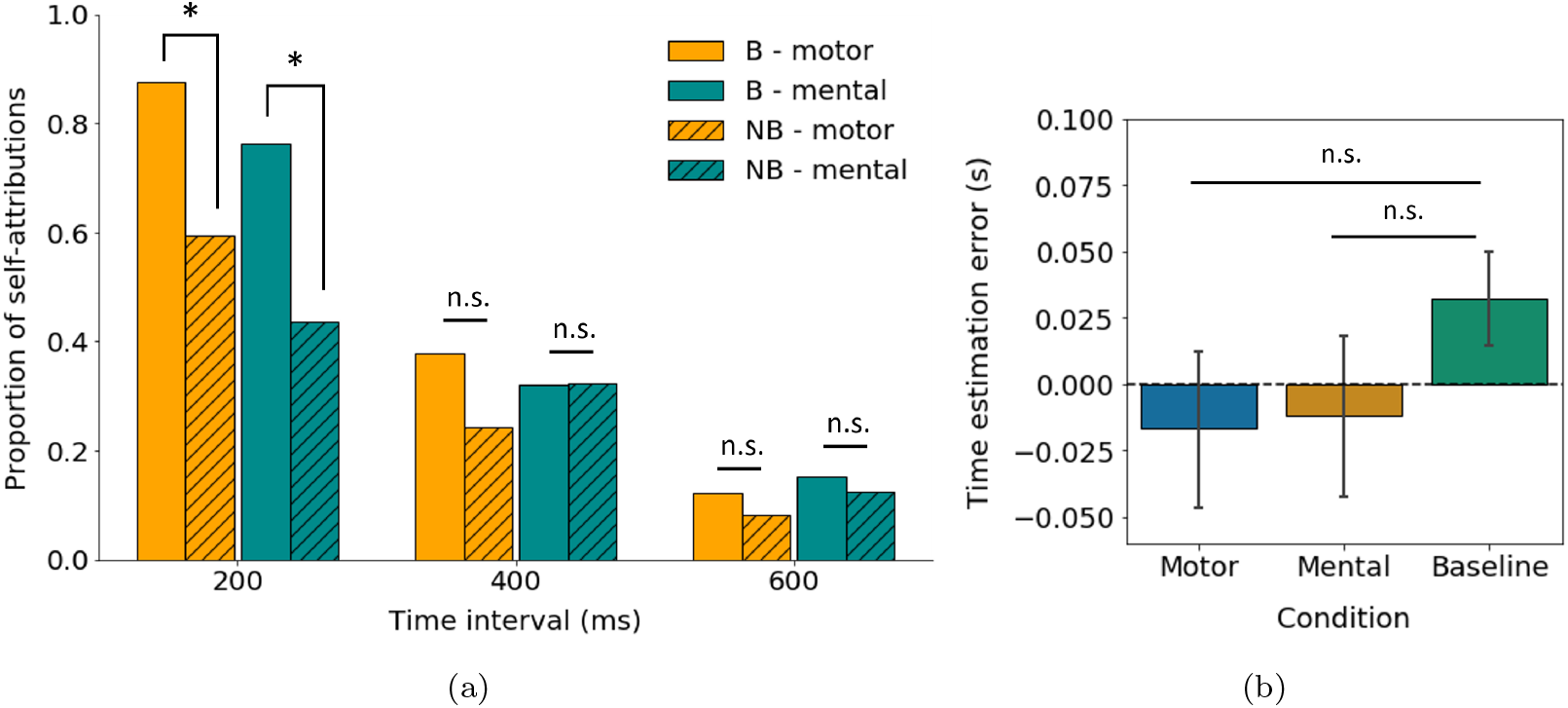
Participants without causal beliefs reported a reduced high-level sense of agency and no intentional binding. (a) Explicit reports: proportion of self-attributions for believers (*B*) and non-believers (*NB*) in each condition and interval. The statistical significance in the comparison between groups is shown for illustrative purposes (the statistical tests have not been carried out on the proportion of self-attributions but on the number of attributions). A significant difference in the number of self-attributions by believers versus non-believers was found for short intervals in both the mental and motor action condition. \**p* < 0.017; ‘n.s.’: non-significant (*p* > 0.017); Chi-Square tests, Bonferroni corrected. (b) Intentional binding: mean error in the estimation of the sound time (with respect to the actual time) for each experimental condition. Errors more negative than the baseline error indicate intentional binding. No significant binding was found for the motor or the mental action condition, indicating the absence of low-level agency. Bars represent 95% confidence intervals. ‘n.s.’: non-significant (*p* > 0.05), paired samples t-test.

We then studied whether intentional binding could be found in the motor and mental action condition for non-believers. For each subject, we compared the mean baseline error with the mean error under the motor or the mental condition. Paired samples t-tests revealed that there was no intentional binding in the motor action condition (*t*(9) = −1.29, *p* = 0.23), nor in the mental action condition (*t*(9) = 0.89, *p* = 0.40). There was also no significant difference between the mean error in the motor and mental action conditions, as shown by an independent samples t-test (*t*(10) = −0.13, *p* = 0.89). Figure 5b illustrates these results. We conclude that the sense of agency was not present at the low level for participants without causal beliefs, as opposed to participants who reported believing that their actions could produce the sound. In this sense, non-believers can be considered as a “control” group, proving that intentional binding is not the mere result of time contiguity, but that at least the belief of causality has to be present.

## 4. Discussion

In the present study, we have investigated the sense of agency arising from motor actions and mental actions in an action-effect paradigm, using implicit and explicit measures of agency.

An important outcome of this study is that we have successfully managed to reconstruct the sense of agency in subjects by using an action-effect paradigm where only beliefs of participants about causality had been manipulated prior to performing the tasks. Indeed, the stimulus was identical in the three experimental conditions: a dot would rotate and after some given time a sound appeared, independent of the participants’ action. However, before each condition, participants were led to believe that the sound could be triggered by a motor action (a key press), by a mental action (thinking about a particular word), or that it would appear spontaneously (baseline condition).

The experimental design was based on the assumption that a sense of agency would be created if prior causal beliefs were sufficiently established. The results of the experiment confirm this assumption: more than two thirds of the total number of participants claimed to be in control of the sound in the motor action block, and more than one third of the subjects thought that their mental action could trigger the sound in the mental action block. The fact that a sense of agency emerged for those participants purely from the manipulation of prior causal beliefs is a critical success of this experiment. This was further confirmed by analyzing the results for subjects who did not have causal beliefs: the sense of agency was reduced or even absent for these participants.

### 4.1. High- and low-level agency

We assessed the high-level JoA by measuring the number of times that a participant would attribute the sound to his/her own action or to the computer, and comparing it for motor and mental actions. We validated the finding that shorter intervals lead to higher agency under motor actions, as confirmed by the higher number of self-attributions for short action-effect delays, and the progressive reduction of self-attributions for medium and large delays. Interestingly, this same pattern was found under mental actions, and no significant difference was found in the number of self-attributions for mental and motor actions, confirming our main hypothesis. This result offers a novel insight to the possibility that a strong SoA arises under non-motor actions, at least at a high level (judgement of agency). Although it has already been suggested by previous studies involving BCIs that a SoA might emerge from mental actions [14, 19], our results show not only that mental actions produce a JoA, but that they do so to a similar extent as motor actions (that is, the number of self-attributions ascribed to mental actions was not significantly different from that of motor actions, in any of the above time intervals); and that the SoA for mental actions disappears progressively under longer action-effect time delays, as occurs for motor actions. Within the framework of the cue integration theory of the SoA [4, 6], these results indicate that in the absence of motor signals, other cues such as prior beliefs or non-motor predictions may contribute to the construction of judgements of agency, compensating for the lack of motor cues. This contradicts other views that propose that motor commands are necessary for a robust sense of agency to emerge, as suggested by accounts based on the comparator model [38].

We also measured the intentional binding over the sound produced by motor and mental actions, as a marker of low-level agency. We replicated the finding that voluntary motor actions produce binding over the effect of those actions: the sound was perceived as occurring earlier than it actually occurred. Critically, we also found that mental actions produced binding over the sound, and that there was no significant difference in the magnitude of the intentional binding effect for motor and for mental actions, once again confirming our hypothesis. This is a major finding of the present study, from which two separate conclusions can be extracted. First, that the intentional binding effect is not to be exclusively linked to motor actions as some authors suggest [39, 34], but is rather a signature of low-level agency for intentional actions in general, as proposed by others [26]. It must be noted that our results are also consistent with the hypothesis put forward by Buehner [28] that the IB effect could reflect pure causality and not intentionality. In any case, both accounts agree in the fact that motor cues are not exclusively required for IB to occur. Second, we conclude that mental actions give rise to a sense of agency not only at a higher conceptual level, but also at a lower perceptual level. This, we claim, is a novel finding in the research on agency. To the best of our knowledge, the current study is the first one to investigate the FoA under non-motor actions.

We contemplate that the construction of the FoA from mental actions presumably relies on general predictive mechanisms, such as counterfactuals, combined with inferential postdictive mechanisms not related to motor cues [6]. In fact, it could be that the FoA for both motor and mental actions is in part based on cognitive predictions independent of motor-based forward models, as suggested by Dogge and colleagues [25]. Based on the pre-activation account of intentional binding proposed by Waszak and colleagues [39], we hypothesize that these general cognitive predictions could pre-activate the perceptual representation of the sensory effect (in this case, the sound), leading to intentional binding (the original proposal by Waszak and colleagues suggested that these predictions were motor predictions).

We also investigated whether a consistent relationship between explicit and implicit measures of agency could be found: if this was the case, the intentional binding effect should be substantially larger for shorter intervals, given that in most short-delay trials participants attributed the authorship of the sound to themselves. Indeed, in the motor action condition we found significant binding for short and medium intervals but no binding for large intervals. This confirms that there is a coherent relationship between the explicit reports of the participants and the implicit measurement of the sense of agency through the intentional binding paradigm. This result is of special relevance because other studies investigating the relationship between explicit and implicit measures of agency have not found a fully consistent relationship between both measures, concluding that they might be driven by dissociable mechanisms [29, 31]. Here instead, we have found that the intentional binding over the sound is a good predictor of the explicit agency reported by the participants. This provides further evidence that the low-level feeling of agency is the major basis for the construction of a higher level judgement of agency, as hypothesized by Synofzik and colleagues [6].

This consistent relationship between implicit and explicit measures also existed for mental actions, in particular for medium and long intervals (binding was found for medium delays but not for long delays). However, the binding over the sound was not significant for short intervals, which was not expected because participants had reported a high amount of self-attributions for short delays in the mental action condition. This result can be interpreted in two alternative ways: either no low-level feeling of agency emerged for non-motor actions under short delays, or a feeling of agency emerged but was not captured by the intentional binding effect. We support the latter option, for the reasons that follow. First, the high-level judgements of agency for 200 ms intervals were higher than for medium intervals. It is widely accepted that the judgements of agency, even if influenced by other factors, are largely based on the low-level feeling of agency. If no FoA arose for 200 ms, but did so for 400 ms, one would expect higher JoA for medium than for short intervals; but this was not the case. Second, the absence of intentional binding for mental actions under short intervals can be explained by the fact that the outcomes of mental actions might be predicted slower than the outcomes of motor actions. Indeed, in our life-long experience interacting with the external environment generally requires bodily movements (BCI are an exception to this). Thus, it is coherent that the predictive mechanisms for the sensory outcomes of motor movements operate at fast timescales (fast arm movements may last less than 200 ms) [40], while the predictions of sensory events triggered by mental actions, which are less common, may be less efficient. In the framework of the pre-activation account of IB [39], this could mean that the predictive mechanisms for mental actions were not fast enough to pre-activate the perceptual representation before the stimulus arrived, and therefore no intentional binding was found. In the absence of an efficient sensory prediction, the FoA might have been constructed through other mechanisms, such as postdictive inferential processes based on the external sensory input and on cognitive priors, as proposed by cue integration theories. Nevertheless, the results of the current study do not allow us to draw a definite conclusion with respect to this matter, and it shall be further investigated whether a feeling of agency may arise for mental actions and short action-effect intervals. For that purpose, other implicit measures such as sensory attenuation paradigms may be useful [41, 42].

In summary, we have found that the sense of agency emerging at the high conceptual level was as strong for non-motor mental actions as for motor actions, and followed a similar pattern for different time intervals. At the low perceptual level, mental actions produced intentional binding in a similar manner as for motor actions, with the exception of short action-effect delays, a matter which will need further investigation. Furthermore, we have found a consistent relationship between the explicit reports and the intentional binding effect, suggesting that this phenomenon might be driven by general predictive mechanisms not limited to motor predictions. Finally, we have found that, in the absence of causal beliefs, the JoA is substantially reduced and the intentional binding effect is not found, suggesting an absence of FoA. This emphasizes the key role of action-effect causality beliefs, which are necessary for the resulting sense of agency.

### 4.2. Limitations and additional considerations

This study presents several challenges and limitations that might be addressed in future work. First, one could argue that the alleged non-motor actions involve motor signals to a certain extent, because participants are told that the software detects facial micro-expressions. Even if participants are told not to produce any voluntary gesture, it is possible that some motor commands are involved in the execution of the action, for example by forcing a ‘static’ facial expression to avoid producing voluntary gestures. Our prediction is that the results would have been very similar had this experiment been carried out with a BCI that did not involve any kind of motor signal; we surmise that any possible contribution of motor cues to the mental action condition is negligible. Another limitation is that many participants did not believe that they were in control in the mental action condition. We think that using a more credible paradigm like an EEG-based BCI would greatly increase the number of participants for whom we could manipulate prior causal beliefs.

A potential limitation of this study is that in usual SoA experiments, participants get to choose when to perform an action, which action to perform or whether to act or not. In our everyday life, the conjunction of these three components (what to do, when to do it and whether to do it or not) gives rise to a strong feeling of intentionality in our actions. The relevance and neuroscientific evidence of the ‘what’, ‘when’ and ‘whether’ components of intentional action have been reviewed elsewhere [43]. In the present study, participants were told which action to perform and when to perform it (and following the instructions did not leave a choice whether or not to perform the action). However, we argue that the decision to follow the instructions was in itself an intention to act that participants executed, and the presence of this intentionality led to a partial version of the ‘what’, ‘when’ and ‘whether’ components. This justifies the fact that participants reported a SoA under our experimental paradigm. We hypothesize that under a real BCI paradigm, where participants could choose when to act or which action to perform, the sense of agency would be stronger than the one measured in this study. This line of research remains an interesting possibility for future work.

It has been recently argued that the sense of agency measured in cognitive neuroscience is ambiguous between different constructs: the sense of agency as a cognitive ability or as a phenomenal character, and bodily versus external constructs of agency. These distinctions yield a total of four distinct constructs [44]. The authors suggest that the different experimental paradigms used to study the SoA are related to different constructs. We propose that the present study focuses on the external construct of agency, given that in the tasks participants were requested to make judgements about an external event. Moreover, our experiment targets both the cognitive ability and the phenomenal character of SoA, since we have used the self-other distinction for the authorship of external outcomes (associated with the SoA as a cognitive ability), and the intentional binding effect, which is related to the phenomenal character of the SoA. In fact, given that we have found a coherent relationship between these measures (self-other distinction and intentional binding), we argue that both constructs of the SoA arise from related mechanisms, and perhaps could be studied interchangeably, at least when referring to the external construct of the SoA.

### 4.3. Implications for BCI research

The findings of this study have important implications for the field of brain-computer interfaces. Previous studies have already presented evidence that subjects performing BCI-mediated tasks report being in control of those actions [14, 19]. The results presented in our study suggest that this sense of agency does not exclusively arise at a pure conceptual level, but also at a low perceptual level, similarly to the SoA for bodily movements. This implies that interaction with BCIs may help reconstruct a strong sense of control, making them promising technologies for the future in a wide range of applications. In particular, the robust feeling of agency evidenced in our study argues in favor of the use of BCIs for the control of external devices, such as prosthetic limbs [45], and for other applications requiring a strong sense of control, such as communication through brain interfaces [46].

Importantly, the high SoA measured for non-motor mental actions indicates that external devices could be controlled in a robust manner not only by BCIs based on motor commands (typically through motor imagery), but also by non-motor-based BCIs, such as steady state visually evoked potential (SSVEP) based BCIs. Nevertheless, it is possible that sensorimotor BCIs provide more robust solutions for applications requiring precise and fast control, as suggested by the absence of intentional binding for mental actions and short delays in this experiment, and as also seen in other studies [19].

Another interesting implication of our study concerns how legal attributions of moral responsibility may have to be rethought in the presence of future BCI technologies. Several authors have already raised issues concerning moral responsibility over BCI-mediated actions: users might be uncertain about their agency in a BCI context, making difficult the attribution of moral (and therefore, legal) responsibility [47, 48]. Moreover, in most legal systems this attribution of responsibility is based on the fact that only “voluntary acts” are punishable, an “act or action” being defined as a “bodily movement” (USA Model Penal Code, Section 2.01). Reflexes and unconscious movements are qualified as involuntary and, thus, are not punishable. However, BCI-mediated actions would not be proper actions under the definition of “act”, even if those are voluntary. The current findings support the idea that BCI-mediated actions should presumably be considered voluntary actions, given that subjects have a sense of agency for mental actions to a comparable extent and execution as they do for motor actions. Thus, we argue that legal systems may need to be adapted to include wider definitions of agency, notably with respect to mentally-triggered external events. The manner in which the notion of agency might be extended to mental actions in the framework of current legal systems is still a matter of debate and will be of critical relevance for the future of BCI technologies in society [47, 49, 48, 18].

### 4.4. Clinical applications

Our experiment suggests that the presence of causal beliefs may induce an illusion of control over external events which are in fact unrelated to the subjects’ actions. Such illusions of control could be used alongside the treatment of psychological disorders typically associated with a loss of control, as is often the case for patients suffering from depression or anxiety [50]. For instance, a recent study has shown that simply activating memories of depression in healthy subjects caused an increase in their estimates of intentional binding compared to activation of baseline memories, thus suggesting a partial loss of sense of agency during depressive episodes [51]. The results of our study suggest that implementing positive causal beliefs during therapy may play an important role in increasing the sense of agency in patients with depression.

Additionally, BCI-based experiments such as the one used in this study might greatly help understand pathological disruptions of the sense of agency, in particular the phenomenon of thought insertion in schizophrenia [52]. Voss and colleagues have explored the mechanisms behind the abnormal experiences of agency exhibited by schizophrenic patients through an IB paradigm [53]. Analogous to that, here we propose that investigating the sense of control over mentally-triggered external events in patients with thought insertion symptomatology may help elucidate mechanisms disrupted under this pathology. It has been suggested that the authorship of thoughts is not a feeling arising at a low level, but rather at a high conceptual level [52]. Thus, one would expect to find a similar intentional binding (low level) for mentally-triggered events in patients suffering from thought insertion compared to healthy subjects; but schizophrenic patients would probably display abnormal attributions of authorship (high level) for the outcomes of these mental actions. This line of research remains to be explored in future work.

## Acknowledgements

RM-B is supported by BFU2017-85936-P and FLAGERA-PCIN-2015-162-C02-02 from MINECO (Spain), the Howard Hughes Medical Institute (HHMI; ref 55008742), The Bial Foundation (grant number 117/18) and ICREA Academia (2016).

